# Pre- and post-oviposition behavioural strategies to protect eggs against extreme winter cold in an insect with maternal care

**DOI:** 10.1101/2021.11.23.469705

**Authors:** Jean-Claude Tourneur, Claire Cole, Jess Vickruck, Simon Dupont, Joël Meunier

## Abstract

Depositing eggs in an area with adequate temperature is often crucial for mothers and their offspring, as the eggs are immobile and therefore cannot avoid exposure to sub-optimal temperatures. However, the importance of temperature on oviposition site selection is less clear when mothers can avoid these potential adverse effects by both moving their eggs after oviposition and providing other forms of egg care. In this study, we addressed this question in the European earwig, an insect in which mothers care for the eggs during several months in winter, frequently moving them during this period. We set up 60 females from two random natural populations (as this species often exhibits population-specific life-history traits and behaviours) under controlled thermal gradients, and recorded the temperature at which they built their nests, tested whether they moved their eggs after an experimental temperature change, and measured the effects on egg development and hatching rate. Our results demonstrate that females indeed select oviposition sites according to temperature, and can move their eggs to reach warmer temperatures. We also show that these warmer temperatures are necessary to ensure egg hatching. Although this set of behavioural thermoregulations is present in the two tested populations, we found a population-specific modality of expression. These included the range of temperatures explored before oviposition, temperature selected at oviposition and dynamics of egg transport following a temperature change. Overall, our study sheds light on a new post-oviposition strategy in female insects that overwinter with their eggs for coping with temperature changes. More generally, it also reveals that egg care and/or egg transport do not prevent behavioural thermoregulation via oviposition site selection and highlights the diversity of behaviours that insects can adopt to enhance their tolerance to global climate change.

## INTRODUCTION

Oviposition site selection shapes the fitness of most oviparous species (Thompson 1988; Refsnider and Janzen 2010; Meunier et al. 2022). This is because choosing the right place to deposit eggs typically provides direct and indirect benefits to egg-laying females, their current eggs, and their future juveniles. This behaviour may first limit the high risk of predation on adult females that is inherent to their lack of mobility during oviposition and favour their direct access to specific food sources necessary for oviposition. For instance, females of the water strider *Aquarius paludum insularis* (Motoschulsky, 1866) avoid ovipositing in sites containing a predator attacking adults only (Hirayama and Kasuya 2013), while females of the orange tip butterfly *Anthocharis cardamines* (Linnaeus, 1758) deposit eggs on plants that have high nutritional value for adults but poor nutritional value for their larvae (Courtney 1981). Oviposition site selection can also provide direct benefits to eggs by limiting the risks of predators finding the eggs or eggs drying out. In the aquatic beetles *Hydroporus incognitus* Sharp, 1869 and *H. nigrita* (Fabricius, 1792), for example, females select waters where no fish can predate on their eggs (Brodin et al. 2006), while in the damselfly *Lestes macrostigmata* (Eversmann, 1836) females prefer to lay their eggs on plants growing in the deeper parts of temporary ponds to minimize the risk of future egg desiccation (Lambret et al. 2018). Finally, oviposition site selection can help future offspring by favouring proximity to suitable habitats and providing direct access to resources necessary for juveniles. This is the case in the sandpaper frog *Lechriodus fletcher* (Boulenger, 1890) and several lady beetle species, where females choose oviposition sites that contain the largest quantities of nutritive resources for their future larvae (Sicsú et al. 2020; Gould et al. 2021).

The effect of temperature on egg development and survival is another potential driver of oviposition site selection by females. Exposure to extreme temperatures can indeed damage living organisms of all ages through alterations in their physiology, immunity and behaviour, which may overall reduce their fitness and/or lead to premature death (Hance et al. 2007; Dillon et al. 2009; Fey et al. 2015; Filazzola et al. 2021). These effects can be particularly strong in eggs because they are immobile and thus unable to escape from environmental temperatures, their shell often provides limited thermal protection, and the development and survival of embryos (contained in eggs) are generally sensitive to subtle changes in surrounding temperatures (Wang et al. 2010; Nicolai et al. 2013; Mortola and Gaonac’h-Lovejoy 2016; Cordero et al. 2018; Yang et al. 2018). As a result, females of many species select oviposition sites according to optimal temperatures for their eggs, such as in the toad-headed agama lizard *Phrynocephalus przewalskii* Strauch, 1876 (Li et al. 2018), the solitary red mason bee *Osmia bicornis* (Linnaeus, 1758) (Ostap-Chec et al. 2021), or the flat-rock spiders *Hemicloea major* (Koch, 1875) (Pike et al. 2012).

By contrast, the importance of temperature for oviposition site selection becomes less clear when mothers can transport their eggs from one location to another, as it may allow them to secondarily adjust the temperature of their eggs throughout development. This transport, included in a broader phenomenon called egg brooding in insects (Machado and Trumbo 2018), is known to allow parents to limit the risk of egg predations or promote egg oxygenation in aquatic species. For instance in the golden egg bug *Phyllomorpha laciniata* (Villers, 1789), females lay their eggs on conspecifics whose mobility improves the avoidance of egg parasitoids (Carrasco and Kaitala 2009). Similarly, in the water bug *Abedus herbeti* Hidalgo, 1935 females lay their eggs on the males’ back, which then move (with the eggs) to ensure that they receive a proper level of oxygenation (Smith 1997). However, whether active egg transport (e.g. parents actively moving their eggs from one location to another) could be an adaptive behaviour by which mothers adjust the thermal needs of the embryo during development remains unexplored. Yet, this process could operate in the European earwig *Forficula auricularia* Linnaeus, 1758. In this complex of cryptic species (Wirth et al. 1998; González-Miguéns et al. 2020) that can be found on almost all continents (Lamb and Wellington 1975; Guillet et al. 2000; Quarrell et al. 2018; Hill et al. 2019), females usually lay their eggs just before or during winter, and then remain with their eggs several weeks or months until they hatch (Lamb 1976; Gingras and Tourneur 2001). During this period, females provide extensive forms of egg care including, for instance, grooming behaviours to remove pathogens, the application of chemical compounds on eggshells to improve resistance against desiccation (Liu et al. 1997; Boos et al. 2014), and fierce protection against predators (Thesing et al. 2015; Van Meyel et al. 2019). Moreover, females are frequently observed transporting their eggs from one location to another by holding them individually between their mouthparts (Diehl and Meunier 2018; Meunier et al. 2020).

Recent results and observations suggest that earwig eggs could benefit from temperature-dependent oviposition site selection and temperature-dependent maternal transport during development. First, the duration of egg development in winter varies from three weeks (e.g. in Southern Europe) to several months (e.g. in North America) (Ratz et al. 2016; Tourneur 2018). This suggests that eggs could be exposed to extremely low (and damaging) temperatures for a very long time if females do not select the oviposition site accordingly and/or do not transport the eggs during development. Second, a recent study shows that prolonging egg exposure to cold (5°C) for 15-day during winter delays the hatching date and development of juveniles to adulthood (which typically takes two months in *F. auricularia*), leads to the production of lighter adult females, and shapes the basal immunity of these females: it increases their overall phenoloxidase activity and reduces the number of haemocytes in females facing a changing social environment (Körner et al. 2018). Because these traits are likely to affect negatively and/or positively the fitness of the resulting adults (Koch and Meunier 2014), temperature-dependent oviposition site selection and egg transport during development could be adaptive strategies in *F. auricularia* mothers. Finally, several laboratory breeding trials indicate that eggs of some populations need to be exposed to near-zero temperatures to trigger embryo development and that subsequently the temperature needs to be increased to continue this development, while others do not (e.g. Wirth et al. 1998; Meunier et al. 2012; Ratz et al. 2016; Tourneur and Meunier 2020).

In this study, we investigated whether European earwig females select an oviposition site based on environmental temperature, and then move their eggs depending on egg age and experimental changes in environmental temperature. Because previous studies point out that the European earwig may show population-specific life-history traits and behaviours (Ratz et al. 2016; Tourneur 2017, 2018; Tourneur and Meunier 2020), we also tested whether temperature-dependent oviposition site selection and egg transport vary between two (randomly selected) populations naturally and experimentally sharing comparable climatic conditions. We set up 60 females from two Canadian populations in two experimental devices (Fig 1) allowing different thermal gradients and then recorded 1) the range of temperatures explored by each female during the 15 days preceding oviposition, 2) the temperature at which females laid their eggs, 3) whether and how mothers transported their clutch along three thermal gradients throughout egg development and finally, 4) how the temperature of these gradients affected juvenile production. Overall, our results reveal that environmental temperature shapes female exploration before oviposition, oviposition site selection, egg transport during development and the production of juveniles. They also show that both oviposition site selection and the dynamic of egg transport during development varies between the tested populations.

**Figure 1.**
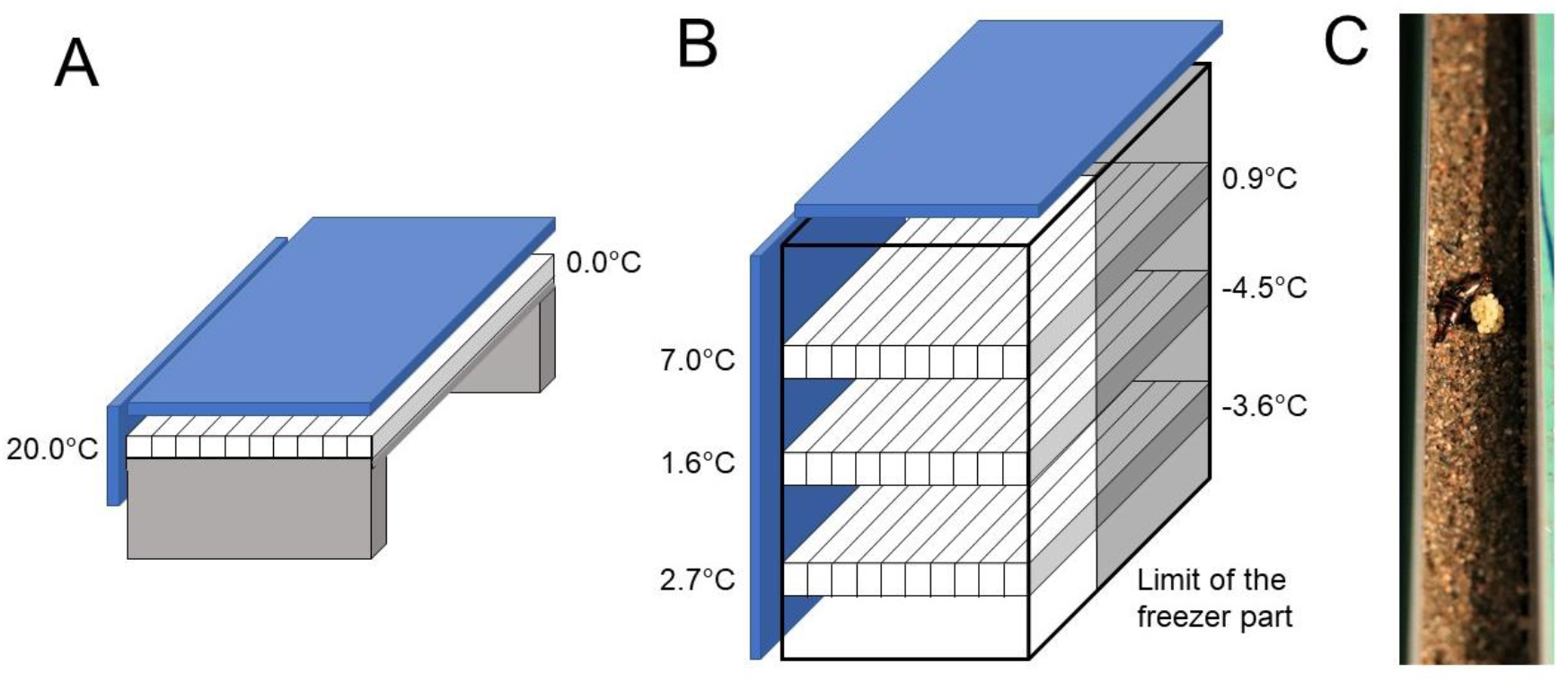
Schema of the thermal setups used in the experiment. (A) Schema of one of the two thermoelectric plates with temperatures linearly ranging from 0°C to 20°C and insulated on every side with thick foam (blue) to ensure complete darkness and thermal isolation. Each plate received 16 rails, each containing one female until they laid eggs and then from the 15th week after oviposition until egg hatching. (B) Schema of the homemade climate cabinet build from a small vertical freezer whose door had been removed and the apparatus covered with plywood enclosure (blue=foam) to ensure complete darkness and thermal isolation, and a vertically sliding front door to allow removal. The cabinet was subdivided into three parts to obtain warm (0.9°C to 7.0°C), cold (−4.5°C to 1.6°C) and intermediate (−3.6°C to 2.7°C) temperature ranges. This cabinet hosted 60 females (20 per temperature range) and their eggs for 15 weeks after oviposition. (C) Picture of a female with its eggs in an experimental rail.

## METHODS

### Earwig sampling and experimental process

The experiment involved a total of 60 *F. auricularia* females field sampled in Harvey station (67°00’52.2”W, 45°38’23.6”N; New Brunswick, Canada; HNB) and St John’s (47°34’42.6”N, 52°44’37.0”W; Newfoundland and Labrador, Canada; SJNL) in September 2020 and then maintained in plastic containers until October 2020. These two populations were selected as independent random units on the basis that they were 2500 km apart by land (1000 km by sea) and had comparable climatic conditions (Fig 2). Subsequent genetic analyses revealed that HNB individuals belonged to the *F. auricularia* genetic clade ‘A’ and SJNL females to the *F. auricularia* genetic clade ‘B’ (see details in Appendix 1 and discussion; Wirth et al. 1998, González-Miguéns et al. 2020). The field sampled females were maintained in groups with males from the same population (sampled at the same time as the females) to allow the completion of the gregarious phase of the life-cycle, which lasts several months during which females mate with multiple partners (Sandrin et al. 2015) and express simple social behaviours (Costa 2006; Weiß et al. 2014). The plastic containers were lined with wet sand, contained two small shelters, and were maintained at room temperature under natural day : light. In October 2020, we isolated each female as this is the period when they usually leave their group to search for a future nesting site (Lamb 1976). To this end, we transferred 36 of 60 females (18 from HNB and 18 from SJNL) to the middle of 36 aluminium rails (Fig 1C; 1.8 x 1. 8 x 72 cm = height x width x length) covered with a layer of wet sand and closed with a plastic cover for subsequent measurement of the temperature range explored before and at oviposition. We isolated the remaining 24 females (12 from HNB and 12 from SJNL) in Petri dishes (diameter 10 cm) covered with moist sand and kept in complete darkness until their use in the measurement of egg transport after oviposition.

**Figure 2.**
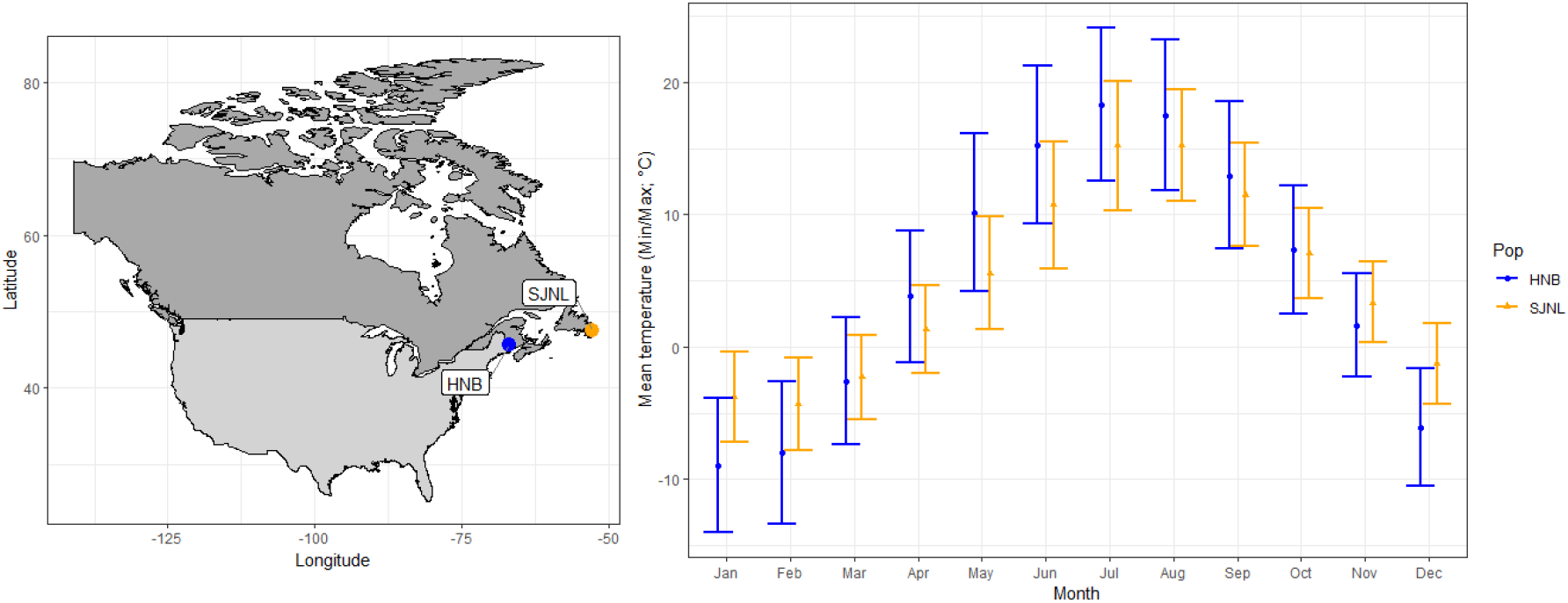
Locations of Harvey Station (HNB, Canada; Blue) and St Johns (SJNL, Canada; Orange), and their monthly variation of temperatures recorded from 1970 to 2000. Dots depict mean monthly temperatures, with whiskers extending from minimum to maximum mean temperatures. These data were extracted from the WorldClim database.

To determine the temperature range explored by each female before egg production and the temperature chosen to deposit eggs, we transferred the 36 aluminium rails containing one female each on thermoelectric plates (AHP-1200CPV-Thermoelectric Cooling America Corporation. 4049W Schubert Avenue. Chicago IL. USA) with temperatures linearly ranging from 0°C to 20°C (Fig 1A and Fig S1 in the Appendix section). We then measured the distance between each female and the coldest edge of its rail every day from the time they were placed in the rails until they produced eggs (including the day of oviposition). Because the date of oviposition greatly varies within populations (up to several months between the first and last oviposition; e.g. Tourneur and Meunier 2020), we standardised our measurement to the temperature range explored by each female 15 days before its own oviposition. To facilitate distance measurement, we divided each rail into 12 zones of 6 cm length and defined the distance between a female and the coldest edge as the centre of the zone she was in. In the very few cases where females were observed between two zones, we assigned females’ location to the colder of the two zones. From field sampling to oviposition, we fed the 36 females twice a week with fresh carrots placed on a soaked cotton pad, plus an artificial diet composed of 1/3 dry power of egg yolk, 1/3 bee collected pollen (Community Apiaries. 576 Plymouth Road, Richmond Corner, New-Brunswick. Canada) and 1/3 cricket powder (Entomo Farms. 31 industrial drive Norwood, Ontario. Canada). We then did not feed females from the day of oviposition to egg hatching, as earwig mothers typically stop foraging during this period (Kölliker 2007).

We then tested whether mothers moved their eggs depending on experimental changes in environmental temperature and/or egg age during the 15 weeks following oviposition. Three days after each female has laid its eggs, we transferred the mother and all its eggs to a new (shorter) aluminium rail (1.8 x 1.8 x 66 cm), which was deposited into a climate cabinet (Fig 1B) providing three non-linear ranges of temperature: warm (0.9°C to 7.0°C), intermediate (−3.6°C to 2.7°C) or cold (−4.5°C to 1.6°C) (Fig 1B and S1). These three thermal gradients encompass the above-ground natural range of temperatures of the two populations during the time females were maintained in our laboratory, i.e. during the natural period of egg care (Gingras and Tourneur 2001; Fig 2). To experimentally change the temperature at which females and eggs were maintained at oviposition, we deposited each female and its eggs in the middle of the new rail (Fig 1C), i.e. at either 5.2°C (warm), 1.2°C (intermediate) or 0.1°C (cold). We then measured the distance (in cm) between the centre of the pile of eggs and the coldest edge of the rail once a week during the 15 following weeks. This measurement of egg transport involved the 36 females used in the measurements of temperature range explored before egg production (see above), and the 24 females (12 from HNB and 12 from SJNL) previously maintained in Petri dishes until oviposition (60 females total). We had not been able to use these 24 females in the electrothermal plates due to the lack of space in the units. Overall, we thus tested 10 HNB and 10 SJNL females in the warm, 10 HNB and 10 SJNL females in the intermediate and 10 HNB and 10 SJNL females in the cold range. In the climate cabinet, the rails were lined with a layer of wet sand and closed with a plastic cover to ensure complete darkness. They were slightly shorter than those used in the thermoelectric plates for reasons of fit.

We finally tested whether the temperatures at which eggs were maintained during the 15 weeks following oviposition affect egg development and hatching rate. Fifteen weeks after oviposition, we transferred the 60 (short) rails containing the mothers and their eggs (i.e. 5 rails per thermal range and population) to thermoelectric plates with temperatures linearly ranging from 0°C to 19°C (and not 20°C due to the shorter rails). This transfer allowed females to have access to a wider thermal range, as expected in Spring – the moment at which eggs typically hatch in *F. auricularia*. We then checked each female daily to record the date of the first egg hatching, the location of the clutch at hatching (based on the distance between the centre of the clutch and the coldest edge of the rail) and the number of newly hatched juveniles. All distance recordings were made very gently so that the females were not disturbed during the observation.

We recorded the thermal gradients present in the different types of aluminium rails to the nearest 0.1°C using 4 channel K type Thermometers (SD. Amazon. CA) connected to four probes located in the sand either at 2.0, 25.5, 49.0 and 72.0 cm (long aluminium rails) or 1.5, 22.5, 43.5 and 64.5 cm (short aluminium rails) of the coldest edge of the rails. The recordings occurred every hour during the entire experiment on either four aluminium rails evenly distributed over the thermoelectric plates or six aluminium rails evenly distributed among the trails and thermal constraints in the climate cabinet. There were no edge effects on temperatures. We then used these recordings to compute linear (thermoelectric plates) and non-linear (climate cabinet) equations linking distance to temperature (Fig S1), which we then used to obtain the temperature of the location of the females and their eggs.

### Statistical analyses

We used a series of eight non-parametric Exact Mann-Whitney Rank Sum tests correcting for tied observations to test the effect of population on the amplitude of temperatures at which females were observed before oviposition, the warmest and coldest temperatures reached by females before oviposition, the location of females at oviposition, the date of oviposition, the number of eggs produced, the number of weeks until egg hatching and the temperature of the area of egg hatching. Changes in the location of the clutch after oviposition were then analysed using parametric Linear Mixed-effects models (LME) in which the week (1 to 15), population (SJNL and HNB), range of temperature (warm, intermediate and cold) and the interaction between these three factors were entered as fixed explanatory factors, while the ID of each female was used as a random factor to correct for multiple measurements. To interpret the significant interaction between the three fixed explanatory factors, we divided the data set according to temperature range (i.e. in three subsets), in which we ran another series of LMEs in which we used week, population and the interaction as fixed explanatory factors, and the female ID as a random factor. When the interaction between weeks and population were significant, we conducted post hoc pairwise comparisons between the temperature of the location where the eggs were initially transferred (i.e. 5.2, 1.18 and 0.11°C) and the temperature of the location where the eggs were observed each week using a series of one sample Exact Mann-Whitney Rank Sum tests. To correct for multiple testing, the P-values of this series of pairwise comparisons were adjusted using the False Discovery Rate (FDR) method (Benjamini and Hochberg 1995). Finally, the likelihood to produce at least one juvenile and the hatching rate were tested using two Generalized linear models with binomial error distribution and corrected for overdispersion. In these models, the population, the range of temperatures and their interaction were entered as explanatory factors, while we used either the presence of at least one hatched juvenile as a binary response variable (1 or 0) or the hatching rate (the number of eggs that hatched divided by the total number of eggs produced) as a continuous response variable. We checked that the assumptions of the LMEs were met using the DHARMa package (Hartig 2020). The analyses were conducted with the software R v4.1.1 (R Core Team, 2017) loaded with the packages *car* (Fox and Weisberg 2019), *exactRankTests* (Hothorn and Hornik 2021), *emmeans* (Lenth 2021) and *DHARMa* (Hartig 2020). The R-script and data are available in an open data repository (see below).

## RESULTS

Earwig females explored non-random ranges of temperature during the 15 days preceding oviposition (Fig 3). The maximum temperature range was greater for the more artic SJNL than the HNB females (median = 12.2 and 7.8°C, respectively; Fig 3A; W = 53, P < 0.001). The warmest temperature at which females were observed was higher in SJNL than HNB females (median = 19.6 and 12.7°C, respectively; Fig 3B; W = 2, P < 0.001), whereas the coldest location was the same for SJNL and HNB females (median = 5.7 and 4.0°C, respectively; Fig 3C; W = 135.5, P = 0.404).

**Figure 3.**
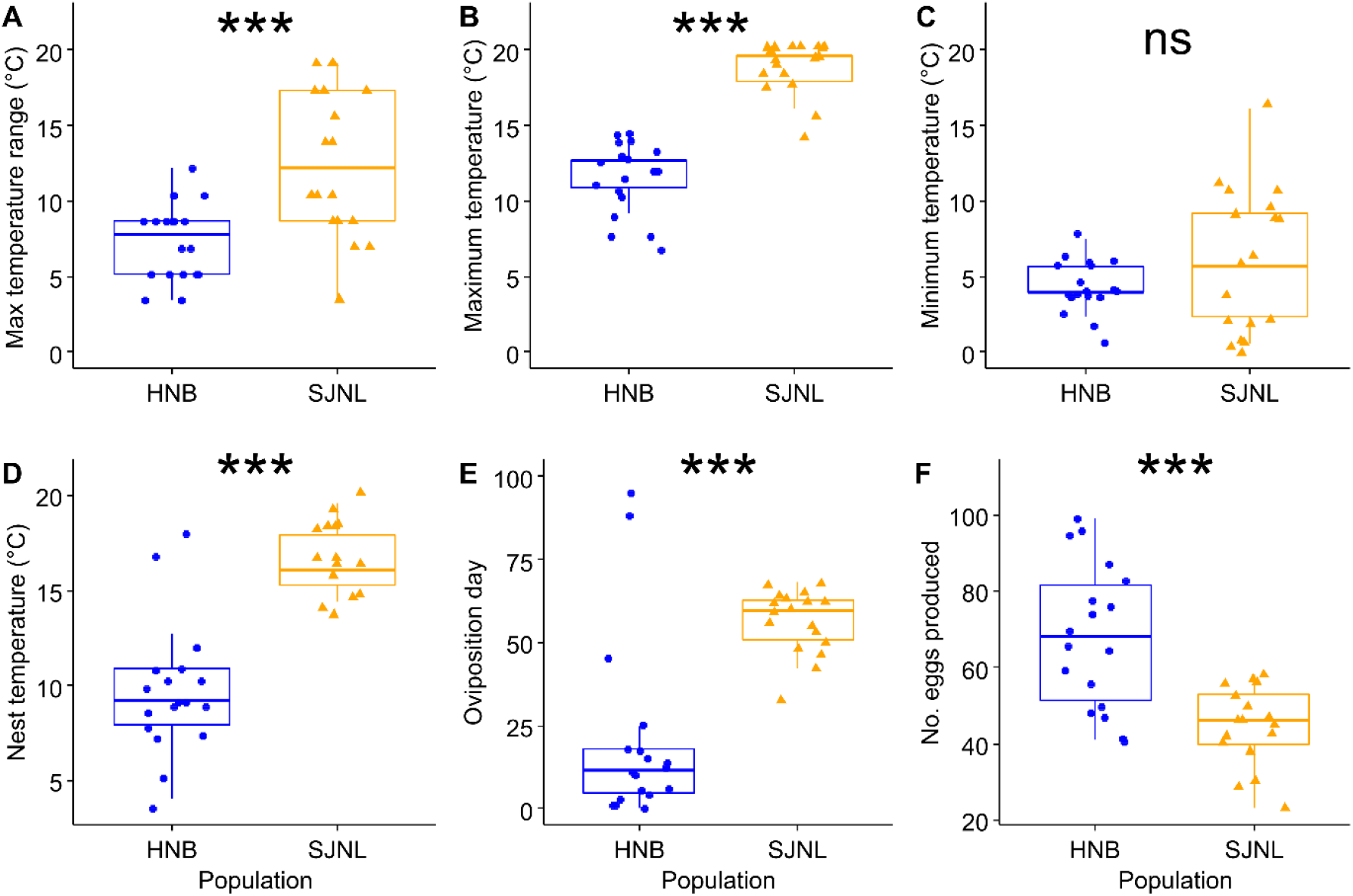
Effect of population on location and reproduction until oviposition. (A) Maximum temperature range, (B) warmest and (C) coldest location at which females were observed during the 15 days preceding oviposition. (D) Location and (E) day at which females laid eggs, and (F) number of eggs produced. Box plots depict median (middle bar) and interquartile range (light bar), with whiskers extending to 1.5 times the interquartile range and dots representing jittered experimental values. ***P<0.001; ns P> 0.05. HNB: Harvey Station. SJNL: St Johns.

Earwig females chose their oviposition site according to the environmental temperature (Fig 3). The temperature of the oviposition site was higher in SJNL compared to HNB females (median = 16.1 and 9.2°C, respectively; Fig 3D; W = 17.5, P < 0.001). Moreover, HNB females produced their eggs earlier in the season (median = 12 and 60 days after first egg production in all females tested, respectively; Fig 3E; W = 38, P < 0.001) and laid overall more eggs (median = 68 and 46, respectively; Fig 3F; W = 261, P < 0.001) than SJNL females.

During the 15 weeks following oviposition, earwig mothers moved their eggs depending on experimental changes in environmental temperature, egg age and population (Fig 4; Triple interaction between weeks, population and range of temperature: Likelihood ratio χ^2_5_^= 33.13, P < 0.001). Egg location depended on an interaction between week and population in the warmest range of temperatures (LR χ^2^= 12.43, P < 0.001), which reveals that SJNL females moved their eggs towards warmer locations about 7 weeks after oviposition, whereas HNB females did not specifically target warmer locations (Fig 4). Egg location also depended on an interaction between week and population in both the intermediate and the coldest range of temperatures (LR χ^2^= 16.53, P < 0.001 and χ^2^= 6.27, P = 0.012, respectively), which shows that both SJNL and HNB females moved their eggs towards warmer locations but that this move started six weeks earlier in SJNL compared to HNB populations (Fig 4). Interestingly, the mothers did not only move their eggs on the sand but built new nests each time they moved their eggs.

**Figure 4.**
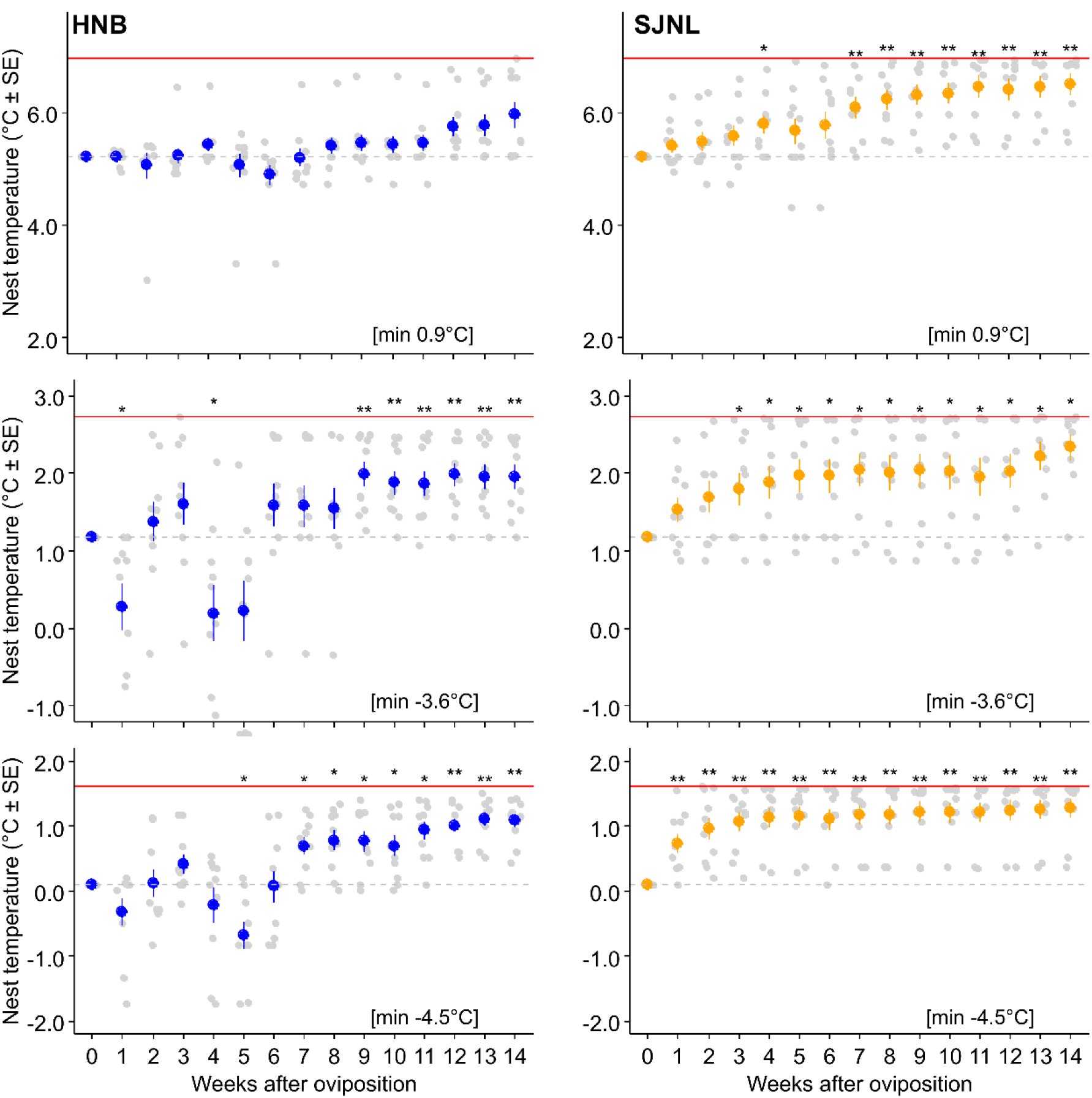
Effect of population and temperature range on egg location during the 15 weeks following oviposition. Brackets indicate thermal ranges. Dashed grey lines indicate the location of mothers and eggs when the experiment was set up. The maximum temperature of each area is indicated with a red line, while its minimum temperature is provided between brackets. Grey dots are jittered experimental values for each clutch of eggs. Coloured dots are mean values ± SE per week. Exact Mann-Whitney tests to compare values of each week to the initial temperature: **P<0.01, *P<0.05. P-values corrected for multiple comparisons. HNB: Harvey Station. SJNL: St Johns.

Finally, the temperatures at which eggs were maintained during the 15 weeks following oviposition affected both egg development and hatching rate. The likelihood to produce juveniles (i.e. that at least one egg hatched) and the egg hatching rate were overall higher for HNB compared to SJNL females (Fig 5A; 53% versus 20%; LR χ^2^= 8.52, P = 0.004 and Fig 5C; 30% versus 12%, LR χ^2^= 6.20, P = 0.013, respectively), overall higher in females previously maintained under the warmest range of temperatures (Fig 5B; 60% versus 25% and 25%; LR χ^2^= 8.09, P = 0.018 and Fig 5D; 36% versus 12% and 15%, LR χ^2^=7.99, P = 0.018, respectively), and not affected by the interaction between these two factors (LR χ^2^= 3.49, P = 0.175 and LR χ^2^=0.61, P = 0.737, respectively). When they hatched, the temperature of the place at which the first juveniles were observed was the same for both populations (Fig 5C; W = 60, P = 0.396), while the eggs took overall more time to develop in HNB compared to SJNL females (Fig 5D; W = 80.5, P = 0.012).

**Figure 5.**
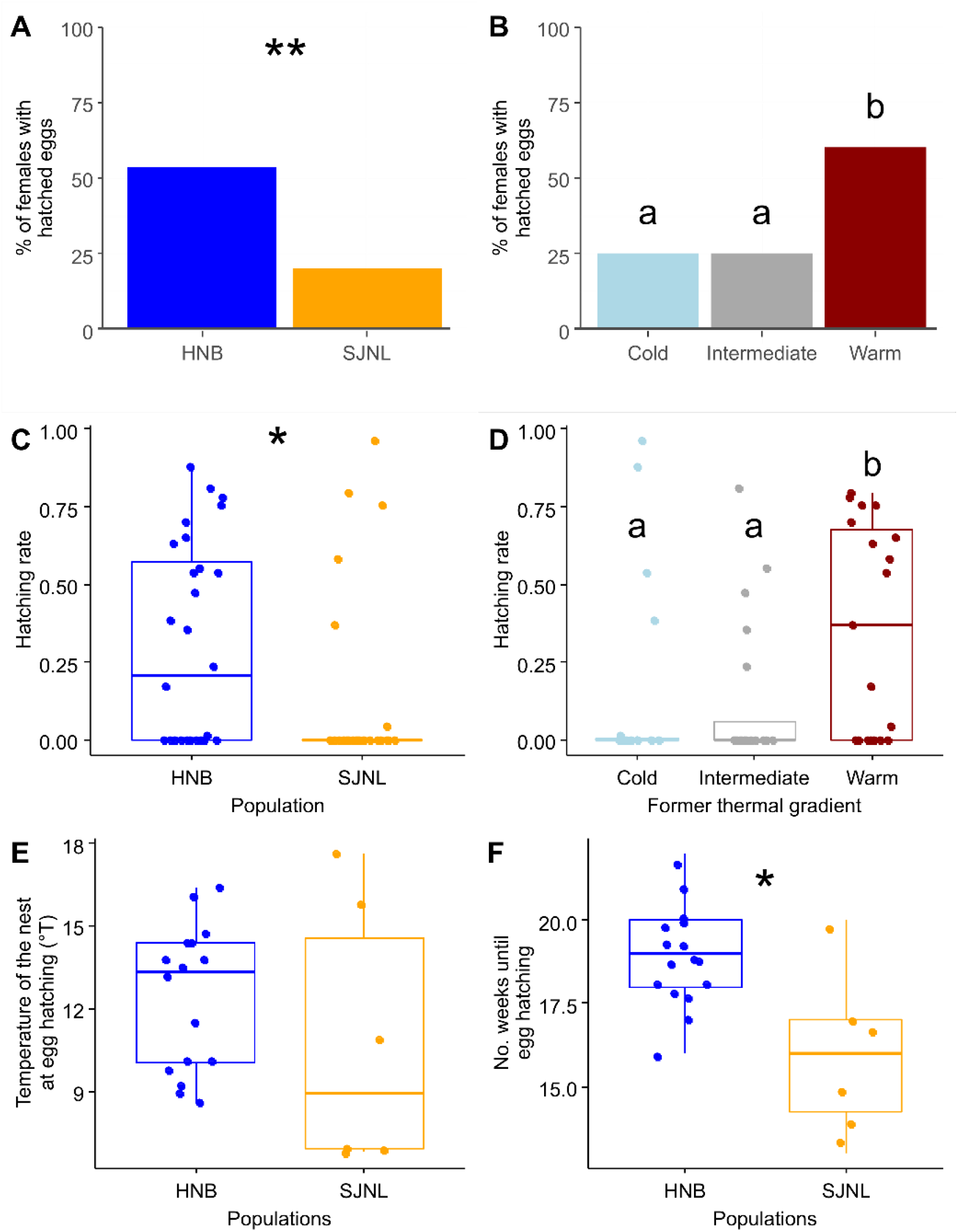
Effect of population and temperature range on (A,B) the percentage of females with at least one hatched egg and (C,D) the egg hatching rate. Effect of population on the location of the eggs at the time of hatching (E) and the number of weeks between oviposition and egg hatching (F). Figures E and F involved females from the warmest range of temperature only, as their numbers were too limited in the other range of temperatures. Box plots depict median (middle bar) and interquartile range (light bar), with whiskers extending to 1.5 times the interquartile range and dots representing jittered experimental values. ns: P>0.05; *P<0.05; **P<0.01;. p = 0.07. Different letters indicate P<0.05. HNB: Harvey Station. SJNL: St Johns.

## DISCUSSION

While temperature often drives oviposition site selection to ensure that the lack of egg mobility does not result in exposure to sub-optimal or extreme temperatures (Pike et al. 2012; Li et al. 2018; Ostap-Chec et al. 2021), the role of temperature on oviposition site selection is less clear when mothers can transport the eggs after oviposition. Here, we show in the European earwig that this capability does not prevent females from selecting oviposition sites according to temperature and that egg transport can indeed help mothers to adjust egg temperature after oviposition. Although this set of behavioural thermoregulations is present in the two tested populations, we found that their modality of expression varied between the two tested populations: St John’s (Newfoundland and Labrador, Canada) females explore a greater range of temperatures before oviposition, lay their eggs in warmer areas, move their eggs quicker toward warm locations when suddenly exposed to cold, but were overall less likely to produce juveniles under our experimental conditions than mothers from Harvey station (New Brunswick, Canada).

Earwig females of both populations chose oviposition sites with temperatures above 10°C, which is in stark contrast to the sub-zero temperatures measured above ground in these two locations during the natural oviposition period (Fig 2). This finding highlights the fact that some ectotherms, such as earwigs, cannot only develop physiological mechanisms to withstand their own freezing during wintering (Toxopeus and Sinclair 2018), but may also prefer to find places exhibiting a large thermal difference from the ground temperature to establish nests and deposit their eggs (Leather et al. 1993). Multiple strategies have been reported in insects to achieve this goal, among which digging nests or burrows to obtain efficient isolation from above-ground temperatures (Davis et al. 2015; Huang et al. 2020), hiding under rocks and nesting into trunks to use the thermal inertia of the substrate as a shelter (Brower et al. 2009; Trájer et al. 2014), and nesting close to human constructions (e.g. underground pipeline, houses, building walls, etc) to benefit from their constant source of heat during winter (Labrie et al. 2008; Trájer et al. 2014). These three strategies are also likely to be adopted by the European earwig, as earwig adults are frequently found in human habitations, underground burrows and under rock and trunks during winter (Goodacre 1997; Gingras and Tourneur 2001; Kölliker and Vancassel 2007; Binns et al. 2021). Moreover, another study suggests that the nest proximity to human constructions could be an effective overwintering strategy for Canadian populations of earwigs (Goodacre 1997).

When experimentally exposed to temperatures below 5.5°C after oviposition, earwig mothers of both populations transported their eggs to warmer locations. Interestingly, these eggs were less likely to hatch when mothers were experimentally prevented from reaching warmer locations, i.e. when mother and eggs were maintained in the cold and intermediate temperature ranges. These results overall support the hypothesis that egg transport is an adaptive post-oviposition behaviour by which earwig mothers protect eggs against extreme cold and/or adjust the thermal needs of their embryos. This discovery sheds light on a new strategy in female insects that overwinter with their eggs for coping with temperature changes (Lee and Dellinger 1991; Sinclair et al. 2003), and emphasizes that the potential costs associated with building a nest and finding a burrow in another location during winter do not prevent such egg transport (Danks 2002). It now calls for future studies on the physiological costs of egg transport for females at a time when they typically stop their foraging activity (Kölliker 2007)(but see Van Meyel and Meunier 2020), and on the impact of temperature variation during egg development (see Figure 4) on hatching success and offspring quality. More generally, the egg transport capability of earwig females combined with their oviposition site selection based on temperature could explain, at least in part, how such an insect with long-overwintering eggs has been able to invade extremely cold climates (Guillet et al. 2000; Quarrell et al. 2018; Hill et al. 2019; Tourneur and Meunier 2020).

Interestingly, St John’s females transported their eggs much earlier than Harvey station females. This suggests that the cold tolerance of eggs in the first few weeks after egg laying is less effective at St John’s than at Harvey station, either due to population-specific egg quality, as reported in numerous oviparous species (Jing and Kang 2003; Stålhandske et al. 2015), or population-specific egg development time, as egg sensitivity to cold often increases when they are closer to hatching (e.g. Gray, 2009). In line with the last scenario, egg development time greatly varies between natural populations of the European earwig (Tourneur and Gingras 1992; Meunier et al. 2012; Ratz et al. 2016; Tourneur and Meunier 2020), and our data show that eggs produced by St John’s females indeed take less time to hatch compared to eggs produced by Harvey station females when reared in similar conditions. At a more general level, it is typically expected that a population-specific cold tolerance of eggs should lead to population-specific timing of oviposition, the less cold-tolerant eggs being laid later in winter than the more cold-tolerant eggs (Tourneur and Meunier 2020). This is again in line with our results: St John’s females laid eggs about one month after Harvey station females. Overall, our results thus demonstrate that winter temperatures are an important factor in the pre- and post-oviposition strategies of earwig females.

Somewhat surprisingly, our study finally reveals that the modality of expression of the reported strategies to protect eggs against severe winter cold varies between the two studied populations. Although our experimental design does not allow us to conclude robustly on the reasons for this variation, we propose three potential explanations. First, this could be due to local adaptation to environmental conditions. In line with this explanation, previous studies reported that multiple traits can vary between populations of the European earwig, such as the number of clutches produced by females, clutch size, juvenile quality, the timing of egg production, duration of egg development and female body mass (Ratz et al. 2016; Tourneur 2017, 2018; Tourneur and Meunier 2020). However, these studies often compared populations with contrasting climatic conditions, which was not the case between St John’s and Harvey stations (Fig 2). Second, the reported variation could be due to population-specific differences in the phenology of females at field sampling. However, this is unlikely to explain our results, as the females were sampled late in the breeding season and our setup control for this potential variation by standardising the measurements around the natural day of oviposition for each female. Finally, the behavioural variation reported between the two tested populations could be due to the presence of different genetic clades. In line with this hypothesis, our genetic analyses reveal that females of Harvey station belong to the genetic clade ‘A’ and females of St John’s to the genetic clade ‘B’ (this was surprising, as this is the first time that the clade ‘B’ is found in Canada outside British Columbia). The European earwig is a complex of cryptic species, for which genetic divergence and reproductive isolation are well established (Wirth et al. 1998; Guillet et al. 2000; González-Miguéns et al. 2020), but the specificity of their life-history traits remains largely unexplored. To date, the only known species-specific trait refers to their reproduction, with females ‘A’ producing one clutch and females ‘B’ producing two clutches (Wirth et al. 1998; Tourneur 2018). However, other studies demonstrate that the number of clutches produced by a female can vary within *F. auricularia* species (Tourneur and Gingras 1992; Ratz et al. 2016) depending on numerous parameters acting during and after the early life of an earwig female (Meunier et al. 2012; Meunier and Kölliker 2012; Wong and Kölliker 2014; Tourneur 2018). If this third hypothesis is true, our results may thus have shed light on the first behavioural difference between ‘A’ and ‘B’ females. Nevertheless, better understanding what drives population-specific dynamics of maternal strategies to protect eggs against cold needs additional studies involving, for instance, several populations of ‘A’ and ‘B’ females and/or population transplants.

Whereas ectotherms typically have very limited to no physiological capacities to limit the costs of exposure to extreme temperatures via internal regulation (Stevenson 1985), some have evolved forms of behavioural thermoregulation to avoid locations with extreme temperatures (Lee Jr. 1991; Terrien 2011). Our study demonstrates that ectotherm females of the European earwig adopt a broad set of behavioural thermoregulations when tending their eggs overwinter. In particular, they select oviposition sites exhibiting specific temperatures and transport their eggs to warmer locations when experimentally exposed to cold. While this set of behavioural thermoregulations is present in the two studied populations, its modality of expression varied between Harvey station and St John’s – either as a result of local adaptation or as species-specific traits among the complex of species composing the European earwig (Wirth et al. 1998; Guillet et al. 2000; González-Miguéns et al. 2020). Overall, our findings emphasize that earwig females have evolved behavioural strategies to mitigate the risks inherent to tending eggs during several months and extreme winter cold. More generally, they highlight the diversity of behaviours that insects can adopt to cope with extreme temperatures, and could favour their tolerance to the effects of moderate climate change.

## DATA ACCESSIBILITY

Data are available online: https://doi.org/10.5281/zenodo.6338078

## SUPPLEMENTARY MATERIAL

Script and codes are available online: https://doi.org/10.5281/zenodo.6338078

## ACKNOWLEDGEMENTS

We warmly thank Wolf U. Blanckenhorn, Nicolas Sauvion, and Anna Cohuet (Peer Community in Zoology referees and recommender, respectively) for their very detailed and constructive comments on this manuscript. Version 3 of this preprint has been peer-reviewed and recommended by Peer Community In Zoology (https://doi.org/10.24072/pci.zool.100012).

## APPENDIX

### Appendix 1 Analyses of the genetic clades of F. auricularia

To determine whether HNB and SJNL females belonged to the same genetic clade (species) of the European earwig (Wirth et al. 1998; González-Miguéns et al. 2020), we analysed the Cytochrome Oxidase I (COI) gene of 6 females per population. This number of females is enough to robustly assess the origin of an entire population, as previous work demonstrated that the two species do not co-exist in the same population (Wirth et al. 1998; Guillet, Guiller, et al. 2000). Genomic DNA was extracted from whole individuals with the NucleoMag^®^ Tissue kit (Macherey Nagel) following manufacturer instructions. The COI gene (658 bp) was amplified from each individual using routine barcoding primers LepF and LepR (Hajibabaei et al. 2006). PCR amplifications was performed with the DreamTaq^®^ PCR Master Mix Kit (Thermo Scientific) using an ESCO Swift Maxi^®^ thermocycler with an initial denaturation step at 95°C (2 min) followed by 35 cycles at 95°C (45 sec), 52°C (60 sec) and 72°C (90 sec) and finally an extension step at 72°C (10 min). PCR products were purified, and Sanger sequenced in both direction using an ABI 3730XL sequencing system (Thermo Fisher Scientific) at Eurofins Genomics Company. Sequences obtained were corrected using Geneious^®^ 9.1.8. The species status of each female was identified from NCBI databases using the BLAST tool (Altschul et al. 1990). The BLAST results were ranked by percent identity and the reference sequences with at least 100% identity to our sequences were used to assign the species status. All sequences obtained in this study have been submitted to GenBank; their accession numbers are OL512959 to OL512964 for the 6 HNB females and OL512965 to OL512970 for the 6 SJNL females. The COI analyses revealed that all (6/6) HNB females belonged to the species “Forficula auricularia A” and all (6/6) SJNL females belonged to the species “Forficula auricularia B”.

**Figure S1.**
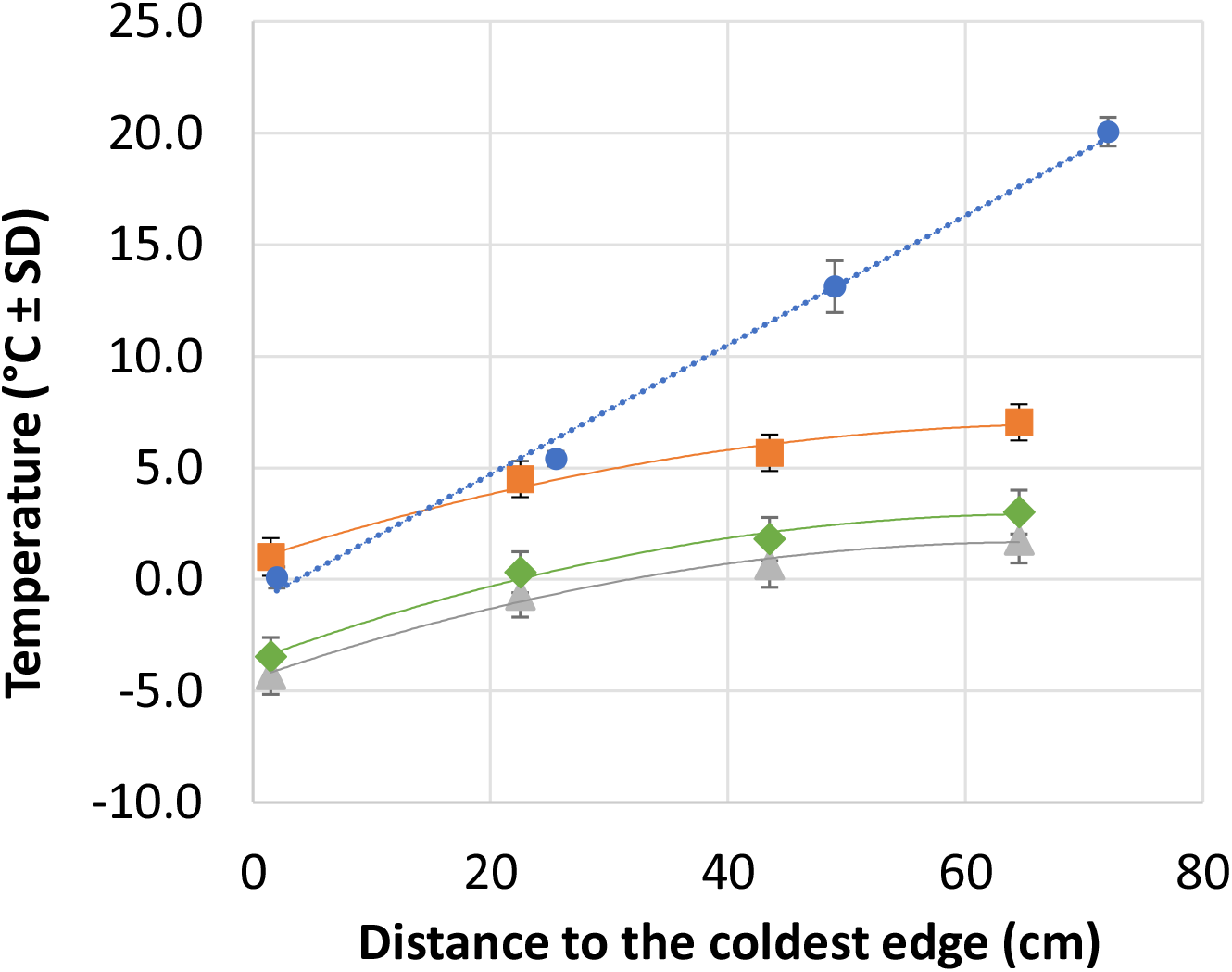
Temperatures measured throughout the experiment in the thermoelectric plates (Blue circles) and climate cabinet (Orange squares, Green diamonds, and Grey triangles) as a function of distance from the coldest edges. In the thermal bridges, the linear regression connecting the measurements is y = 0.2889x - 0.3452. In the climate cabinet, the polynomial regressions connecting the measurements differed between the warmest (Orange square; y = −0.0012×2 + 0.1714x + 0.8747), intermediate (Blue circle; y = - 0.0015×2 + 0.1955x - 3.643) and coldest (Grey triangles; y = −0.0014×2 + 0.1842x - 4.445) temperature ranges.

